# Mechano-adaptation: Exercise-driven Piezo1 & Piezo2 augmentation and chondroprotection in articular cartilage

**DOI:** 10.1101/2024.08.02.606183

**Authors:** Xingyu Jing, Alexander Kotelsky, Yaxin Zhang, Robert Dirksen, Sandeep Mannava, Mark Buckley, Whasil Lee

## Abstract

Chondrocytes in adult joints are mechanosensitive post-mitotic quiescent cells with robustly expressed both Piezo1 and Piezo2 ion channels. Here, we examined the mechano-adaptation and Piezo modulations in articular chondrocytes using a mouse exercise model. We first found differential expression patterns of PIEZO1 and PIEZO2 in articular chondrocytes of healthy knee joints; chondrocytes in tibial cartilage (T) exhibit significantly higher PIEZO1 and PIEZO2 than femoral chondrocytes (F). Interestingly, a few weeks of exercise caused both PIEZO1 and PIEZO2 augmentation in F and T compared to the sedentary control group. Despite the increased expression levels of these mechanosensors, chondrocytes in exercised cartilage exhibit significantly reduced mechanical susceptibility against 1mJ impact. PIEZO1 modulation was relatively more rapid than PIEZO2 channels post-exercise. We tested the exercise-induced effect using Piezo1-conditional knockout (Pz1-cKO; Agc1^CreERT2^;Piezo1^fl/fl^). Pz1-cKO mice exhibit diminished exercise-driven chondroprotection against 1mJ impact, suggesting essential roles of Piezo1-mediated mechanotransduction for physiologic-induced cartilage matrix homeostasis. In addition, using a mouse OA model, we further found the modulated PIEZO1 in chondrocytes, consistent with reports in Ren et al., but without PIEZO2 modulations over OA progression. In summary, our data reveal the distinctly tuned Piezo1 and Piezo2 channels in chondrocytes post-exercise and post-injury, in turn modulating the mechanical susceptibility of chondrocytes. We postulate that Piezo1 is a tightly-regulated **biphasic biomarker**; Piezo1 antagonism may increase cellular survival post-injury and Piezo1 (with Piezo2) agonism to promote cartilage ECM restoration.

## Main Text

Cellular mechanosensation and mechanotransduction are crucial for tissue homeostasis and survival in all living organisms. Piezo1 and Piezo2 have been confirmed to display distinct expression patterns and gating kinetics in response to mechanical stimuli, where the gain- or loss-of-function mutations alter the energetics of Piezo1 or Piezo2, resulting in the imbalanced intracellular ion homeostasis and associated with hundreds of human channelopathies [1, 2]. Distinct gating and expression patterns contribute to diverse cellular mechanotransduction pathways in tissue-specific manners. Understanding cellular mechanosensitivity *in situ* or *in vivo* over diverse mechanical loading environments is crucial to improving Piezo-associated channelopathy.

Mechanical cues heavily influence cartilage tissue homeostasis, and articular chondrocytes in skeletally matured load-bearing cartilage are post-mitotic cells maintaining the cartilage extracellular matrix (ECM) [3-6]. Articular chondrocytes are intrinsically mechanosensitive with robustly expressed mechanosensitive Ca^2+^-permeating ion channels, such as Piezo1, Piezo2, and TRPV4. We and others have reported the essential roles of Piezo1 and Piezo2 sensing hyper-physiologic loading *in vitro* [7, 8]. Piezo1 has been suggested as a potential therapeutic target for osteoarthritis (OA) due to the inflammatory cytokine-induced augmentation, also due to the Piezo1-dependent chondrocyte death [7-10]. In contrast to OA-associated abnormal mechanotransduction, physiologic loading-driven ECM remodeling has been understood dominantly via TRPV4-mediated mechanotransduction *in vitro* and *in vivo* [11-23]. However, Piezo1 and Piezo2-mediated chondrocyte mechanotransduction under physiologic loading conditions still needs to be explored.

In this study, we directly explored the gene regulations of Piezo1 and Piezo2 and the mechanical susceptibilities of chondrocytes using exercise and osteoarthritic mouse models. To examine the physiologic mechanical stimuli-driven effect on chondrocytes, mice were housed with the Voluntary Wheel Running (VWR) apparatus (Med Associates Inc.) and examined joints post-exercise [24-26]. To condition the abnormal osteoarthritic environment, mice underwent a non-invasive anterior cruciate ligament injury (ACL-I), the most common joint injury increasing early onset OA [27-30]. We observed the modulated of Piezo1 and Piezo2 channels post-exercise and post-injury by immunohistochemistry and the mechanical susceptibility of chondrocytes *ex vivo* using a custom-built mechanical impact device [30-32]. This report provides the evidence of the exercise-driven Piezo1 and Piezo2 modulations and essential roles of Piezo1-mediated chondrocyte mechanotransduction in ECM remodeling and chondro-protection. These localized Piezo modulations may exist in other mechanosensing tissues, including heart and lung, and our data alert the need on understanding the spatiotemporal mechano-adaptation post-physiologic and pathophysiologic stimuli.

## Result

### The higher PIEZO1 and PIEZO2 in femoral and tibial chondrocytes in knee joints of wildtype mice

We first examined the cartilage thickness and expression patterns of PIEZO1 and PIEZO2 channels of control mice (C57BL6/J female 3-5-month-old mice) by histochemistry. The articular cartilage thickness of mouse joints is similar between femoral and tibial cartilage (∼100 μm); yet, the thickness of hyaline cartilage (HC, uncalcified zone above tidemark) is more significant in tibia (T) than femur (F) (∼24 μm vs 50 μm). (**Fig.1A-C**). Interestingly, both PIEZO1 and PIEZO2 channels are robustly expressed in chondrocytes in HC but not in calcified cartilage (CC) (**Fig.1d**). Furthermore, both channels are more robustly expressed in tibial chondrocytes than femoral chondrocytes (**Fig.1e-f**). Tibial cartilage is known to endure relatively continuous and higher mechanical stress than femoral cartilage [33-35]; these differential expression levels of PIEZO1 and PIEZO2 may suggest the cellular mechano-adaptation to cope with the local mechanical environment in different compartments of cartilage.

**Fig 1.**
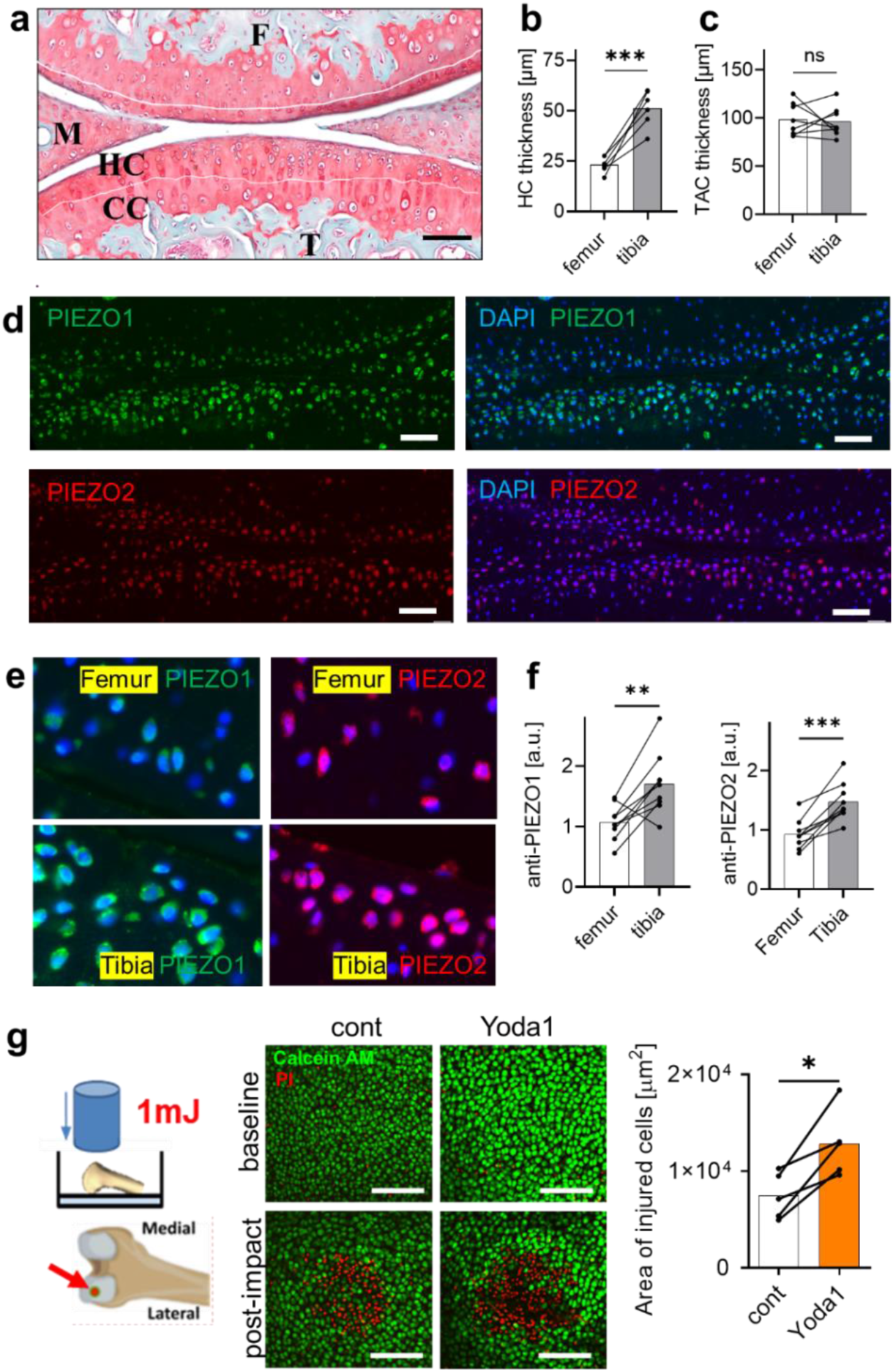
Distinct expression levels of PIEZO1 and PIEZO2 channels in femoral and tibial articular cartilage in C57BL6 mice (3-5 month-old). (a) A representative Saf-O-stained micrograph. (b) Hyalin cartilage (HC) and total articular cartilage (TAC) thickness of femur and tibia; bar = 100μm, F=femur, T=tibia, M=meniscus. (d-e) Representative IF images of anti-PIEZO1 (green), anti-PIEZO2 (red) antibodies, DAPI (blue). Bar = 50μm. (f) Robust anti-PIEZO1 and PIEZO2 intensity in HC. The higher expression of both PIEZO1 and PIEZO2 in tibial chondrocytes than femoral chondrocytes (N=5-7). (g) Schematic diagram of mechano-death assay (left) and representative images (control vs. +Yoda1) of pre- and post-1mj impact with live/dead (green/red). Increased injured area post-injury in Yoda1-treated cartilage than contralateral cartilage. Paired student t-test, *p<0.05; **p<0.001; ***p<0.0001.

### PIEZO1 activation by Yoda1 increases chondrocytes’ mechanical susceptibility

To confirm the roles of Piezo1 on mechanical susceptibility of chondrocytes *in situ*, we pre-treated femoral cartilage of hindlimb with Yoda1, a Piezo1-specific chemical agonist (Piezo2-specific agonists have yet to be discovered). We then quantified the cellular susceptibility against mechanical impact on the femoral head of control mice (female C57BL6/J) *ex vivo*, as described in [30, 31]. The Yoda1-treated group showed a significantly increased area of chondrocyte death compared to the untreated contralateral limbs (**Fig.1g**). This novel finding not only confirms the functional expression of PIEZO1 channels in articular chondrocytes but also underscores their role in modulating chondrocyte survival against mechanical injury, thereby adding a new dimension to this area of research.

### Exercise-driven PIEZO1 and PIEZO2 augmentation in articular chondrocytes

To measure the effect of physiologic loading on chondrocytes, we examined mouse knee joint samples of sedentary and exercised mice groups. Mice were individually housed in a VWR cage for 1-week and 2-week. Sedentary group (Sed, locked wheels), 1-week exercised group (Exer_1w), 2-week exercised group (Exer_2w) ran 0, ∼55, and ∼152 km, respectively (**Fig.2b, Table S1**). We quantified expression levels of the key mechanosensing ion channels of chondrocytes - PIEZO1, PIEZO2, and TRPV4 – by immunohistochemistry. Interestingly, exercise gradually increased the expression levels of PIEZO1 and PIEZO2 in chondrocytes compared to sedentary group (**Fig.2c-h**). The PIEZO1 and PIEZO2 augmentation was more pronounced in tibial chondrocytes than femoral chondrocytes. In addition, the PIEZO2-null chondrocytes decreased over exercise in both femoral and tibial cartilage (**Fig.S1c-d**). Contrast to PIEZO1 and PIEZO2, TRPV4 expression was decreased in femoral chondrocytes and was constant in tibial chondrocytes post-exercise; as well the TRPV4-null chondrocytes showed increasing trend over exercise in both femoral and tibial cartilage (**Fig.2i-k, Fig. S2e-f**). In summary, these immunohistochemistry data reveal that exercise-driven mechanical cues modulate Piezo1 and Piezo2 mechanosensors, indicating the mechano-adaptation of chondrocytes to their mechanical environments. The upregulated PIEZO1 and PIEZO2 may play substantial roles in chondrocyte mechanotransduction and cartilage remodeling, which was understood mainly with TRPV4-mediated chondrocyte mechanotransduction.

**Fig 2.**
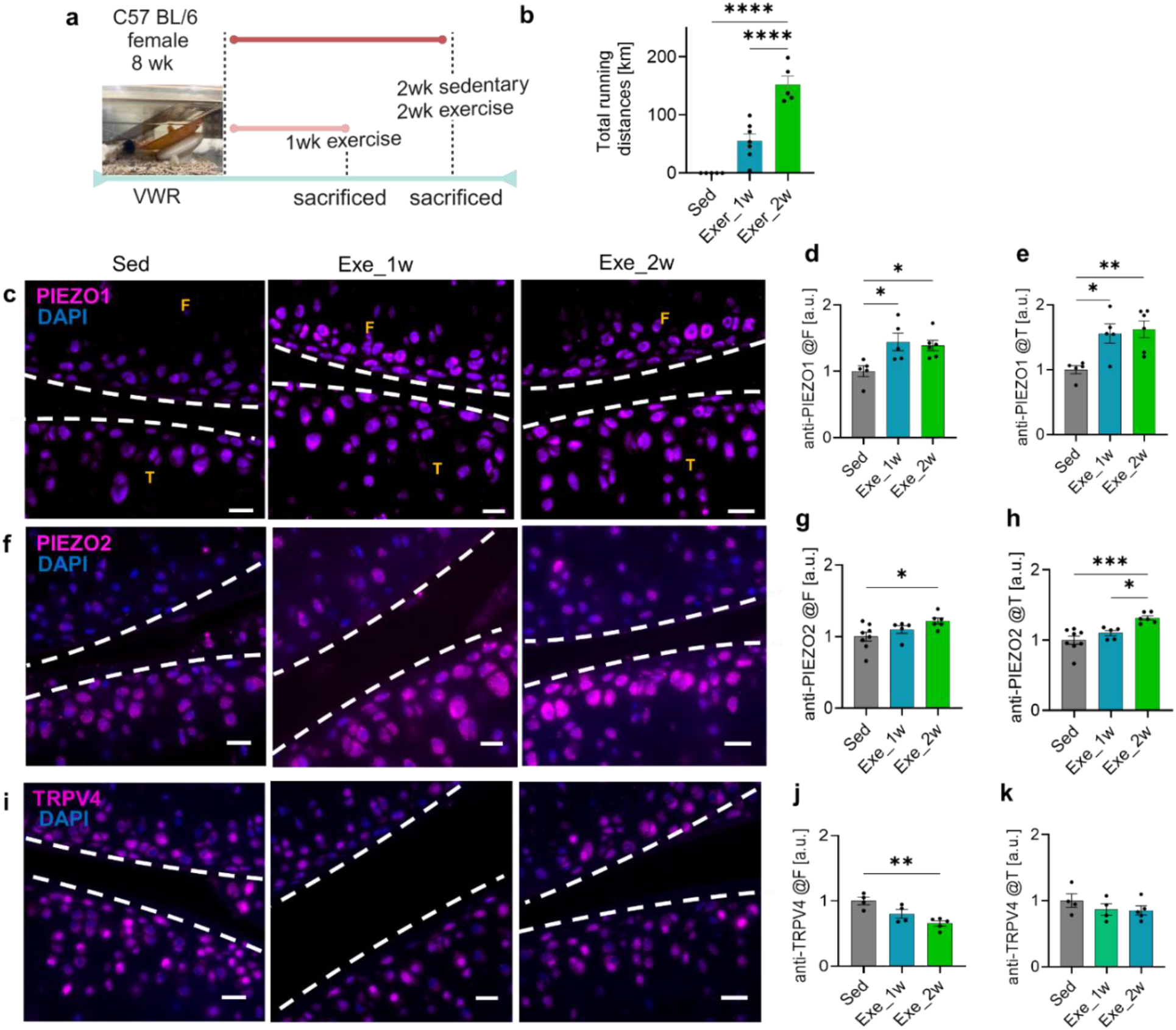
Upregulated PIEZO1 and PIEZO2 post-exercise. (a) Schematic timeline for VWR. (b) Total running distance of each group. (c,f,i) Representative anti-PIEZO1, anti-PIEZO2, and anti-TRPV4 immunofluorescent images of sedentary, 1wk exercise, and 2wk exercise groups. Bar=20μm; F=femur; T=tibia (d, e, g, h) Increased PIEZO1 and PIEZO2 expression in femoral and tibial chondrocytes post-exercise, (j, k) but not TRPV4. Quantified by ImageJ and QuPath. N=5-8/group. Mean±SEM; one-way ANOVA with Tukey’s multiple comparisons. *p<0.05, **p<0.001, ***p<0.0001.

### The sedentary group exhibits similar levels of PIEZO1 in femoral and tibial cartilage

To find the sedentary mice (reduced movement with locked VWR) induce PIEZO modulation, we examined the expression levels of PIEZOs in cells of three compartments of sedentary mice: femoral cartilage, tibial cartilage, and meniscus. Contrast to wildtype mice shown in **Fig.1f**, sedentary mice exhibit reduced PIEZO1 levels in tibial cartilage (**Fig.3a,c**). Meanwhile, PIEZO2 expression continues to be higher in tibial chondrocytes versus femoral chondrocytes, like the wildtype controls (**Fig.3a-d**). Notably, PIEZO1 expression was evident in almost all chondrocytes in HC (95%), while PIEZO2+ cells were 75∼95% in femoral and tibial HC (**Fig.3e-f**). These reduced expression patterns of PIEZO1 and PIEZO2 suggest a potential cellular mechano-adaptation to the reduced mechanical environments.

**Fig 3.**
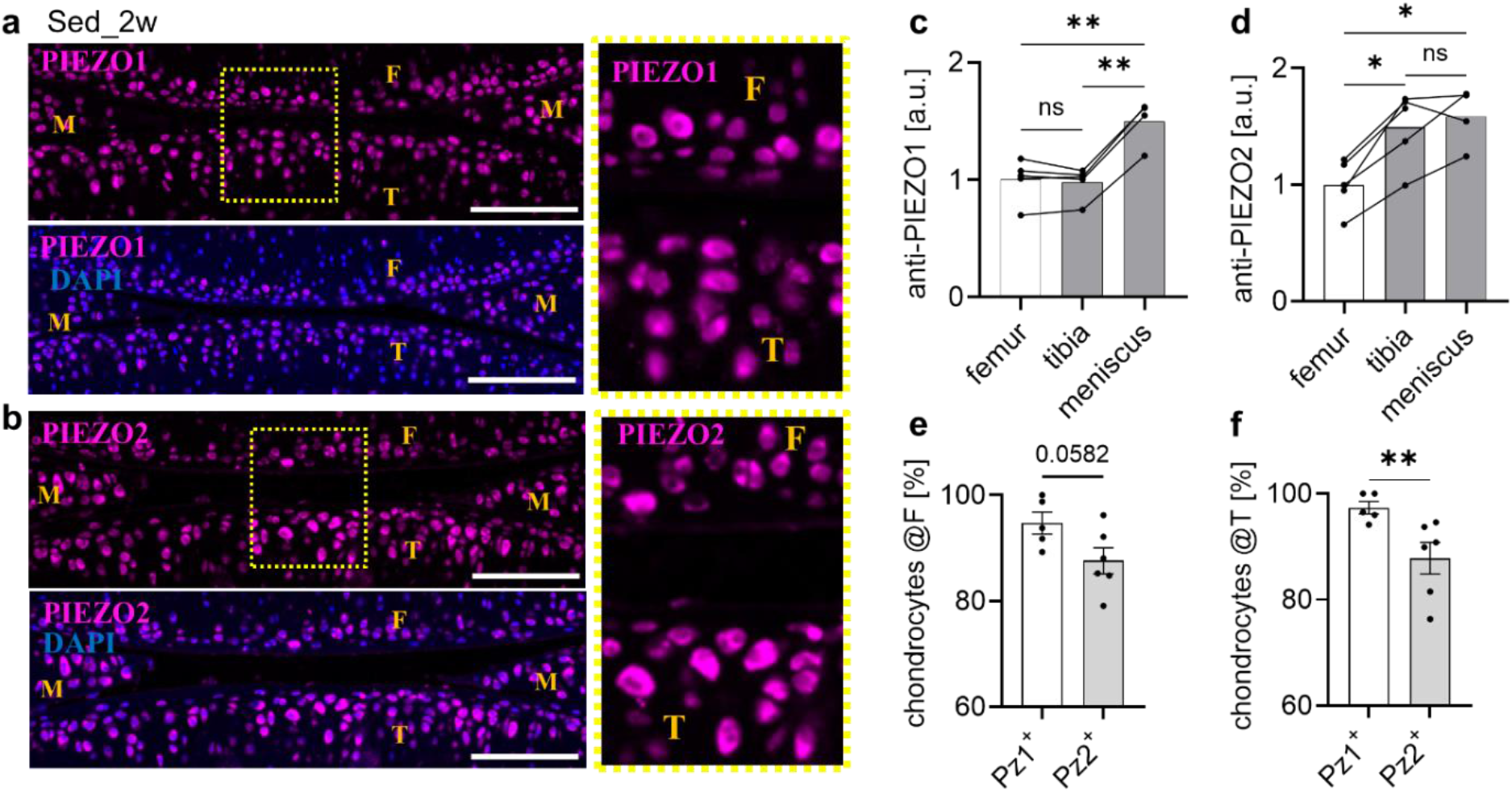
Mice in sedentary (locked VWR, 2 weeks) group exhibit the modulated PIEZO1 and PIEZO2 in femoral and tibial chondrocytes. (a, b) Representative IF images of anti-PIEZO1 (magenta) and anti-PIEZO2 (magenta), DAPI (blue). Bar = 50μm. (c) Anti-PIEZO1 intensity is similar between femoral and tibial chondrocytes, both significantly lower than meniscus. (d) Anti-PIEZO2 intensity is continued to be higher in tibial chondrocytes than femoral chondrocytes. One-way ANOVA. *p<0.05, **p<0.01. (e-f) PIEZO1^+^ chondrocytes are ∼95% in femoral and tibial cartilage, yet PIEZO2^+^ chondrocytes are reduced to ∼85% in femoral and tibial cartilage of sedentary mice. N=4-7 mice/group. Student t-test, **p<0.01.

### Exercised cartilage exhibits reduced mechanical susceptibility

To identify if the augmented PIEZO1 and PIEZO2 channels contribute to the higher injurious loading-induced chondrocyte death *in situ*, we harvested knee joints of sedentary and exercised mice and measured the mechanical vulnerability of chondrocytes against mechanical impact (1mJ) *ex vivo*, as performed in **Fig.1g**. Despite the augmented PIEZO channels, exercised groups exhibit significantly attenuated areas of chondrocyte death compared to sedentary controls, the 4-week exercised group showed almost complete chondroprotection against mechanical injury (**Fig.4a-b**). This exercise-induced chondroprotective effect cannot be explained by the sole hyperphysiologic-strain sensing roles of Piezo1 and Piezo2, which were identified by us and other laboratories by *in vitro* isolated chondrocyte assays [7, 8]. Instead, this effect can be interpreted by the exercise-induced chondrocyte mechanotransduction and biosynthesis for cartilage matrix remodeling. As a proof-of-concept, we compared cartilage thickness and Perlecan levels, one of the key PCM molecules with relatively faster turnover [36, 37], between the sedentary and exercised groups. We found that the HC thickness and the total cartilage thickness were indifferent between sedentary and exercised groups (2 weeks) (**Fig.4c-d**).

**Fig 4.**
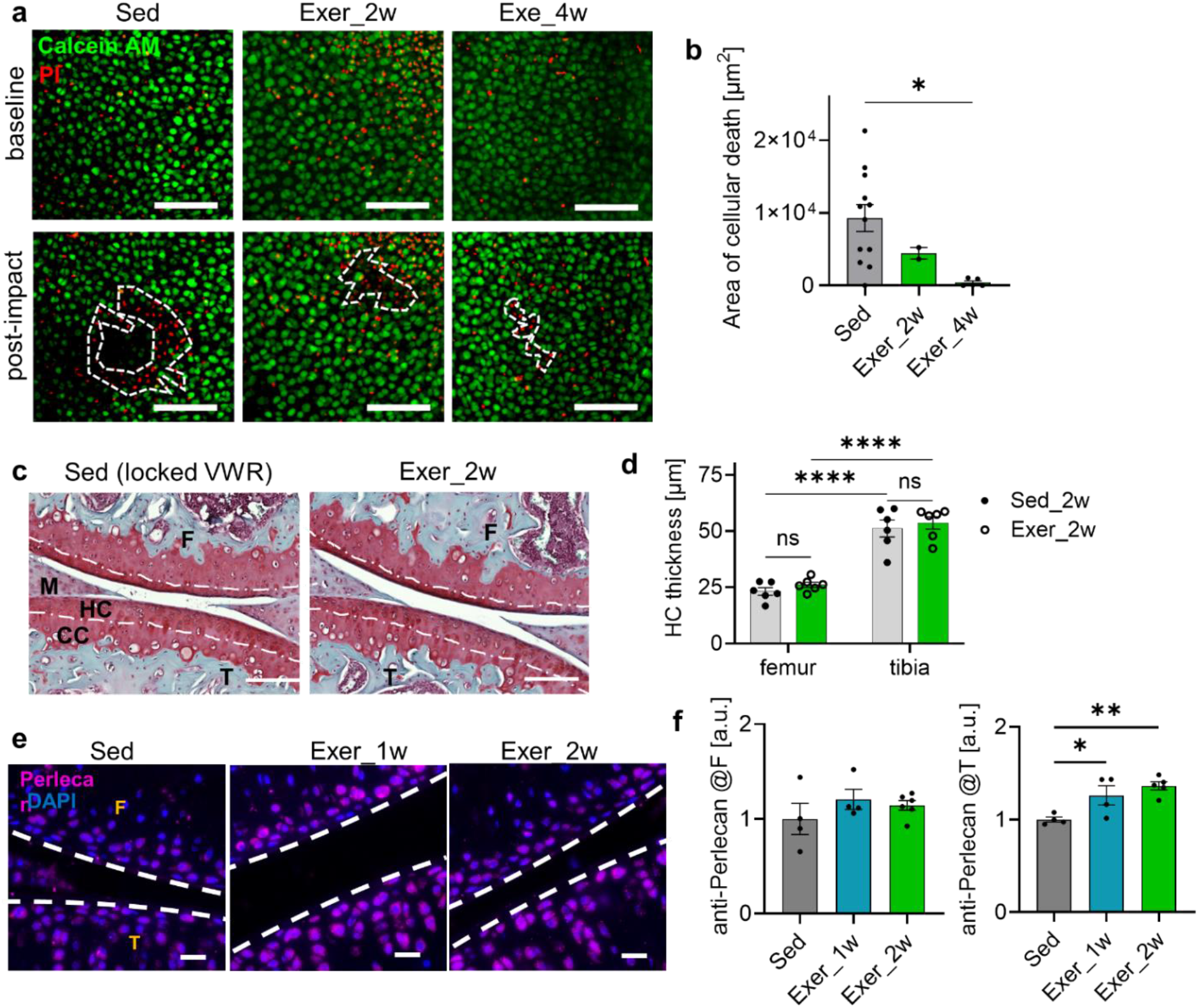
Chondroprotection against impact post-exercise. (a) A representative Safranin-O-stained micrograph of sedentary and 2wk exercise groups. (b) The thickness of HC (hyaline cartilage) was unchanged by 2 weeks of exercise. Bar=100μm. Two-way ANOVA, ****p<0.0001. (c-d) Exercised cartilage were mechanically more resilient with the decreased area of the 1mJ impact-induced chondrocyte death. Calcein (green), PI (red), Bar=100μm. (e) Representative anti-Perlecan IF images of sedentary, 1wk exe, and 2wk exe groups. Perlecan (magenta), DAPI (blue) Pronounced anti-Perlecan intensity in tibial chondrocytes than femoral chondrocytes post-2wk exercise. Unpaired t-test, *p<0.05, **p<0.01.

Notably, the increased Perlecan level was detectable in the tibial cartilage of the 1-week exercise group, and the 2-week exercise group showed more statistically increased anti-Perlecan (**Fig.4e-f**). This anti-Perlecan increase was not statistically significant in femoral chondrocytes; more prolonged exercise effects or another sensitive assay would detect the exercise-driven matrix remodeling in femoral cartilage.

### Articular cartilage of Piezo1-cKO mice suppressed the exercise-induced chondroprotection against mechanical injury

To verify the role of Piezo1 in chondrocytes’ susceptibility *in situ*, we performed the mechano-death assay using the cartilage-specific Piezo1-cKO mice (Agc1^CreERT2^;Piezo1^fl/fl^; Pz1-cKO). We first examined Piezo1 mRNA levels by taking the humoral cartilage of mice at 12 weeks (pulled four mice/sample), and confirmed partial knock-down of Piezo1 in Pz1-cKO cartilage than littermate controls (**Fig.5a-b**). Next, we quantified the area of chondrocyte death using a mechano-death assay. The Pz1-cKO group exhibits a significantly larger area of chondrocyte death compared to the Pz1^fl/fl^ littermate controls (**Fig.4c-d**). To assess the effect of reduced Piezo1 in exercise-driven chondroprotection, we compared the chondrocyte susceptibility of Pz1^flfl^ and Pz1-cKO post-4 weeks of exercise. As shown in **Fig.5e-f**, 4-weeks of exercise did not improve chondrocyte vulnerability against mechanical impact. These data collectively indicate the essential roles of Piezo1 in the mechanical susceptibility of chondrocytes, potentially due to the cartilage matrix degeneration, even with the exercise.

**Fig 5.**
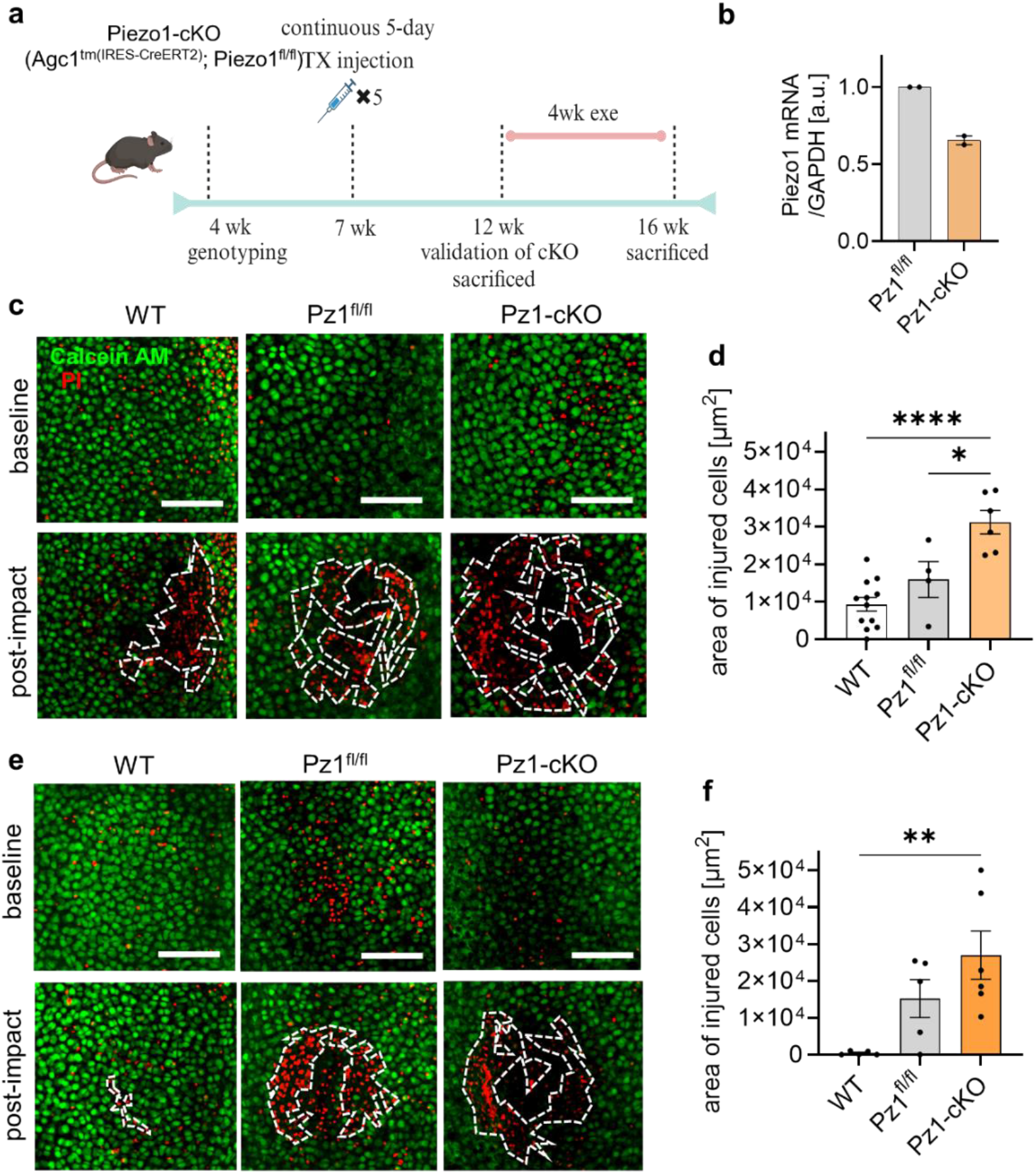
Deleted exercise-driven chondroprotection in Piezo1-cKO (Agc1^CreERT2^;Piezo1^fl/fl^) mice. (a) Timeline of tamoxifen injection and exercise of cartilage-specific Piezo1-cKO mice. (b) Partially decreased Piezo1 mRNA level in femoral head chondrocytes of Piezo1^fl/fl^ and Piezo1-cKO mice. Total RNA was extracted from the femoral head of mouse hip cartilage, 4 mice were pulled in each sample. (c) Representative live/dead micrographs pre- and post-impact. (d) Significantly larger area of dead chondrocytes in Piezo1-cKO mice than littermate Piezo1^fl/fl^ mice or wildtype control. (e) Representative live/dead micrographs pre- and post-impact in 4 wk exercised groups. (f) Significantly increased area of dead chondrocytes in Piezo1-cKO mice than littermate Piezo1^fl/fl^ and wildtype controls. Scale bar=20μm. N = 4∼8 mice/group. One-way ANOVA; *p<0.05; **p< 0.01; ****p<0.0001

### In OA, PIEZO1 is the dominantly modulated mechanosensory in chondrocytes, not PIEZO2

Recent studies by Ren et al. and Wang et al. showed the Piezo1 augmentation in mouse and rat OA cartilage post-trauma, as well as the reduced OA degree with GsMTx4-treatment, pan-Piezo inhibiting peptides [38, 39]. The modulation of Piezo2 in chondrocytes and the mechanical susceptibility of chondrocytes are yet to be explored in the context of OA. This time, we used the anterior cruciate ligament injury (ACL-I)-induced mouse post-traumatic osteoarthritis (PT-OA) with saline or GsMTx4-treatment (8-16week old female C57BL6/J mice) weekly [30, 40]. ACL-I-induced mouse OA model where the ACL-injury is one of the most common joint traumas causing post-traumatic OA. We previously reported that the 8-week-old female mice with a bilateral ACL-I showed OA characteristics, including cartilage degeneration and the increased the mechanical vulnerability of chondrocytes at the age of 12 weeks old [30]. Using the Saf-O/FG-stained micrographs, we found alleviated cartilage degradation and OA degrees in the GsMTx4-treated group than the saline-treated group, consistent with [38, 39] (**Fig.6b-c**). Next, we quantified areas of chondrocyte death by 1 mJ impacts and confirmed the significantly reduced cell death area in the GsMTx4-treated group than the saline-treated group (**Fig.6d-e**).

**Fig 6.**
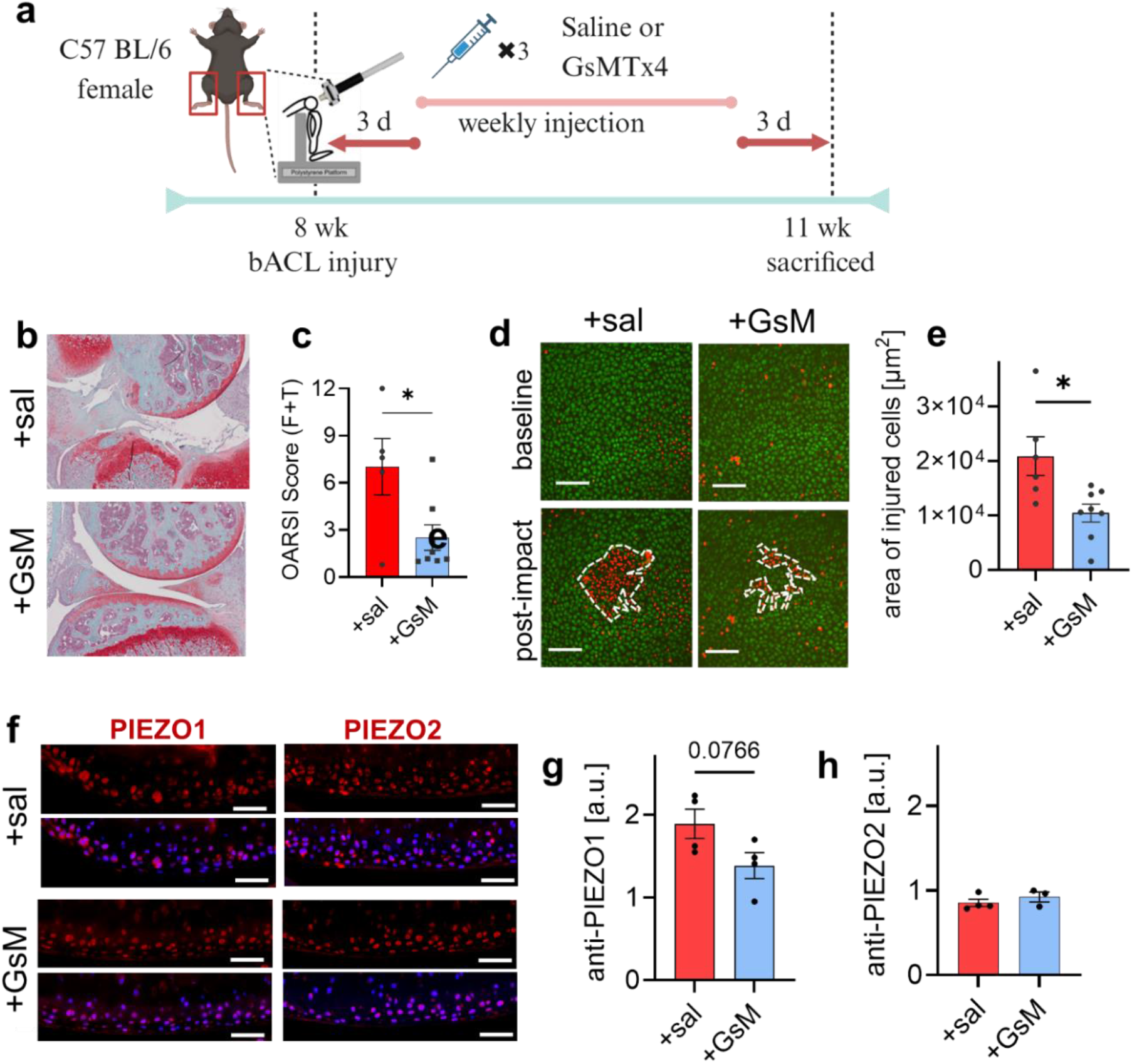
PIEZO1 modulation in femoral cartilage of ACL-injury induced OA mouse with saline or GsMTx4 treatments; yet unchanged PIEZO2 expression levels. (a) Schematic diagram of bilateral ACL-injury (ACLI)-induced mouse OA model and the timeline of saline or GsMTx4 treatment. (b-c) Representative Saf-O-stained micrographs of knee joints of saline or GsMTx4 treated mice post-ACLI. Significantly reduced OARSI score (added femoral and tibial scores) in GsMTx4-treated group than saline-treated group; consistent with Ren et al. (d-e) Representative live/dead (green/red) images of pre- and post-1mj impact. Significantly smaller injured area in GsMTx4-treated group than saline-treated groups. (f) Representative anti-PIEZO1(red) and anti-PIEZO2 (red) micrographs of femoral cartilage in saline- and GsMTx4-treated groups. (g) PIEZO1 expression is higher in saline-treated group than GsMTx4-treated group; consistent with Ren et al. (h) Unchanged PIEZO2 levels in saline- and GsMTx4-treated groups. (g) Unpaired t-test; *p<0.05. Comparison to uninjured control mice is in **Fig**.

We further quantified the PIEZO1 and PIEZO2 expression levels by immunohistochemistry and found PIEZO1 augmentation in OA cartilage of the saline-treated group as compared to the uninjured control or the GsMTx4-treated injured groups (**Fig. 6f-h, Fig.S2**). However, anti-PIEZO2 levels were consistent between the uninjured control group, the saline-treated injured group, and the GsMTx4-treated injured group. These data suggest distinct regulation of Piezo1 and Piezo2 in exercised or osteoarthritic cartilage. It is important to understand Piezo1-mediated mechanotransduction and Piezo2-mediated mechanotransduction for cartilage regeneration and degeneration.

## Discussion

Piezo1 and Piezo2 are evolutionarily conserved calcium-permeating mechanosensitive (MA) channels with 42% homology [1]. Although they share similarity, Piezo1 and Piezo2 have different expression patterns and are associated with diverse types of human channelopathy [1, 41]. Chondrocytes in load-bearing joints are highly sensitive to mechanical stimuli and express both Piezo1 and Piezo2 robustly [8, 10, 42]. In this study, we directly compared the expression patterns of Piezo1 and Piezo2, their exercise-driven modulation, and the mechanical susceptibility of chondrocytes *ex vivo* using mouse models.

We observed the non-uniformly expressed Piezo1 and Piezo2 channels in different regions of murine knee articular cartilage. Chondrocytes in the hyaline cartilage (HC) show more robust expression of PIEZO1 and PIEZO2 channels compared to chondrocytes in the calcified cartilage (CC) in mice. Additionally, chondrocytes in the tibial HC exhibit higher levels of PIEZO1 and PIEZO2 compared to those in the femoral HC. When subjected to exercise-induced loading, both femoral and tibial chondrocytes show increased Piezo1 and Piezo2 channels, but not TRPV4. On the other hand, the sedentary group displays reduced levels of PIEZO1 and PIEZO2 in HC chondrocytes. These findings indicate that regions experiencing higher mechanical loading demonstrate increased expression of Piezo1 and Piezo2, indicating the mechano-adaptation of chondrocytes through Piezo1 and Piezo2 modulation. Notably, these Piezo modulations are distinct from those seen in chondrocytes associated with osteoarthritis, which only show an increase in Piezo1 but not Piezo2. While TRPV4 is well-characterized as a mechanosensing channel in loading-related cartilage biomechanics [5, 21, 43, 44], our data suggest that mechano-adaptation involves modulated Piezo1, Piezo2, and TRPV4 channels, highlighting the essential roles of PIEZO channels following physical activities either independently or synergistically with TRPV4.

Chondrocyte survival and mechanical susceptibility can indicate cartilage resilience and OA progression. Several *in vitro* studies have shown the roles of Piezo1 and Piezo2 as injurious loading sensors that lead to chondrocyte apoptosis. To understand the effects of increased Piezo1 and Piezo2 on chondrocyte death, we quantified the mechanical susceptibility of chondrocytes *in situ* and *ex vivo* post-exercise. Despite the elevated PIEZO1 and PIEZO2 post-exercise, chondrocytes were significantly resilient against mechanical impact. Mechanical trauma-induced cellular death depends on various factors and intracellular and extracellular molecules involved in mechanotransduction. In Piezo1-cKO mice, the exercise-induced chondroprotection was eliminated, suggesting potential roles of Piezo1 in cartilage matrix remodeling. The combined roles of Piezo1 and Piezo2 may play critical roles in matrix remodeling, as OA cartilage with increased Piezo1 alone showed a degenerated extracellular matrix.

Articular chondrocytes are surrounded by a pericellular matrix (PCM) which forms a structure unit known as a chondron. The PCM serves to protect chondrocytes and helps to transmit cartilage deformation in response to mechanical stimuli [45]. When cartilage is compressed, chondrons in HC undergo significant volume changes, and the PCM experiences more deformation due to its relatively lower modulus [36]. Perlecan is one of the key PCM molecules that mediates communication within the matrix by interacting with Col6, and this interaction has been suggested to have a protective effect on the cartilage [36, 37]. In exercised cartilage, tibial chondrocytes show significantly increased levels of perlecan along with augmented PIEZO1 and PIEZO2, but not TRPV4. Nims et al. recently reported that transcriptomic RNAseq data revealed not only catabolic but also anabolic effects in 3D cultured chondrocytes following Yoda1 treatment under unloaded conditions [46]. The enhanced synergy of PIEZO1 and PIEZO2 post-exercise would induce PCM remodeling to cope with or withstand cyclic mechanical stress. Understanding the interplay among cell mechanics, ECM/PCM mechanics, and PIEZO mechanosensors will emerge as a crucial focus of future research in elucidating the causes and treatments of diseases in mechanosensitive tissues, including load-bearing cartilage disorders.

In summary, our data shows the exercise-driven dual Piezo1 and Piezo2 augmentation in adult quiescent post-mitotic chondrocytes in articular cartilage. It appears that the chondrocytes adapt to mechanical stress by regulating the Piezo1 and Piezo2 channels. In OA, only Piezo1 is augmented, and chondrocytes more susceptible to mechanical stress. We propose that Piezo1 is a tightly-regulated biphasic biomarker and can be a therapeutic target for diagnosing and treating cartilage disorders. Future research is needed to delineate how Piezo1 and Piezo2 contribute to the protective effects of exercise on cartilage and the metabolic pathways involved *in situ* and *in vivo*. Moreover, these localized modulations of Piezo channels may also occur in other mechanosensitive tissues. Understanding the transient or localized Piezo-mediated mechano-adaptation may offer valuable insights into therapeutic strategies targeting Piezo channels for various human diseases.

## Acknowledgements

The authors thank Dr. Thomas Suchyna for continuous research discussion and for the GsMTx4. We also thank Center for Musculoskeletal Research (CMSR) Histology Core and University of Rochester Committee on Animal Resources (UCAR) for their technical assistance. We also thank Sukhee Lee, Brian Wise and Paromita Kundu working on the Piezo1 knockout mouse line.

## Author contributions

WL designed the study. XJ, AK, and YZ performed experiments. XJ, AK, YZ, RD, SM, MB, and WL analyzed and interpreted the data. XJ, AK and WL participated in drafting the manuscript for important intellectual content.

## Data and materials availability

All data are available in the manuscript or the supplementary materials.

## Supplementary Materials

Materials and Methods

Figs. S1 to S4

Table S1

